# Establishment of a human three-dimensional chip-based chondro-synovial co-culture joint model for reciprocal cross-talk studies in arthritis research

**DOI:** 10.1101/2021.02.19.431936

**Authors:** Mario Rothbauer, Ruth A. Byrne, Silvia Schobesberger, Isabel Olmos Calvo, Anita Fischer, Eva I. Reihs, Sarah Spitz, Barbara Bachmann, Florian Sevelda, Johannes Holinka, Wolfgang Holnthoner, Heinz Redl, Stefan Tögel, Reinhard Windhager, Hans P. Kiener, Peter Ertl

## Abstract

Rheumatoid arthritis is characterised by a progressive, intermittent inflammation at the synovial membrane, which ultimately leads to the destruction of the synovial joint. The synovial membrane, which is the joint capsule’s inner layer, is lined with fibroblast-like synoviocytes that are the key player supporting persistent arthritis leading to bone erosion and cartilage destruction. While microfluidic models that model molecular aspects of bone erosion between bone-derived cells and synoviocytes have been established, RA’s synovial-chondral axis has yet not been realised using a microfluidic 3D model based on human patient in vitro cultures. Consequently, we established a chip-based three-dimensional tissue co-culture model that simulates the reciprocal cross-talk between individual synovial and chondral organoids. We now demonstrate that chondral organoids, when co-cultivated with synovial organoids, induce a higher degree of cartilage physiology and architecture and show differential cytokine response compared to their respective monocultures highlighting the importance of reciprocal tissue-level cross-talk in the modelling of arthritic diseases.

## Introduction

In today’s ageing society disorders affecting the human musculoskeletal apparatus, also referred to as musculoskeletal diseases (MSD) are of great clinical and scientific interest. In particular, arthritis describes various pathophysiological conditions, including inflammatory arthritis (IA) or osteoarthritis (OA). Inflammatory arthritis (IA) types including rheumatoid arthritis (RA) are in contrast to OA systemic inflammatory autoimmune conditions often found in diarthrodial joints ^1^. Overall, arthritic diseases frequently lead to swelling, pain and impaired movement of joints, severely affecting the quality of life.^2,3^ Fibroblast-like synoviocytes (FLS) are the primary synovial cell type studied in RA.^4^ In healthy synovia, FLS lubricate the joint (e.g. lubricin or hyaluronan), which ensures homeostasis and function. FLS are also responsible for synovial lining formation and the maintenance of the inner joint membrane (Fig 1A, left panel).^5^ In RA, the synovium is the main joint tissue structure infiltrated at the synovial sublining layer by immune cells and is a critical player in osteoclast-mediated bone erosion and pannus formation as well as enzymatic cartilage degradation (Fig 1A. right panel).^6,7^ During the onset and progression of RA, FLS are stimulated by cytokines secreted from activated and migrating immune cells such as macrophages, B cells and T cells^8^ (e.g. IL-1β, TNFα, IL-6, IL-17), and in turn also secrete cytokines (e.g. IL-6, IL-15, IL-23), chemokines (e.g. IL-8. CXCL12) and degradative enzymes (various metalloproteases as well as cathepsins).^8–15^ In this context, IL-6 is a central pro-inflammatory molecule promoting joint inflammation and destruction.^16,17^

**Fig. 1.**
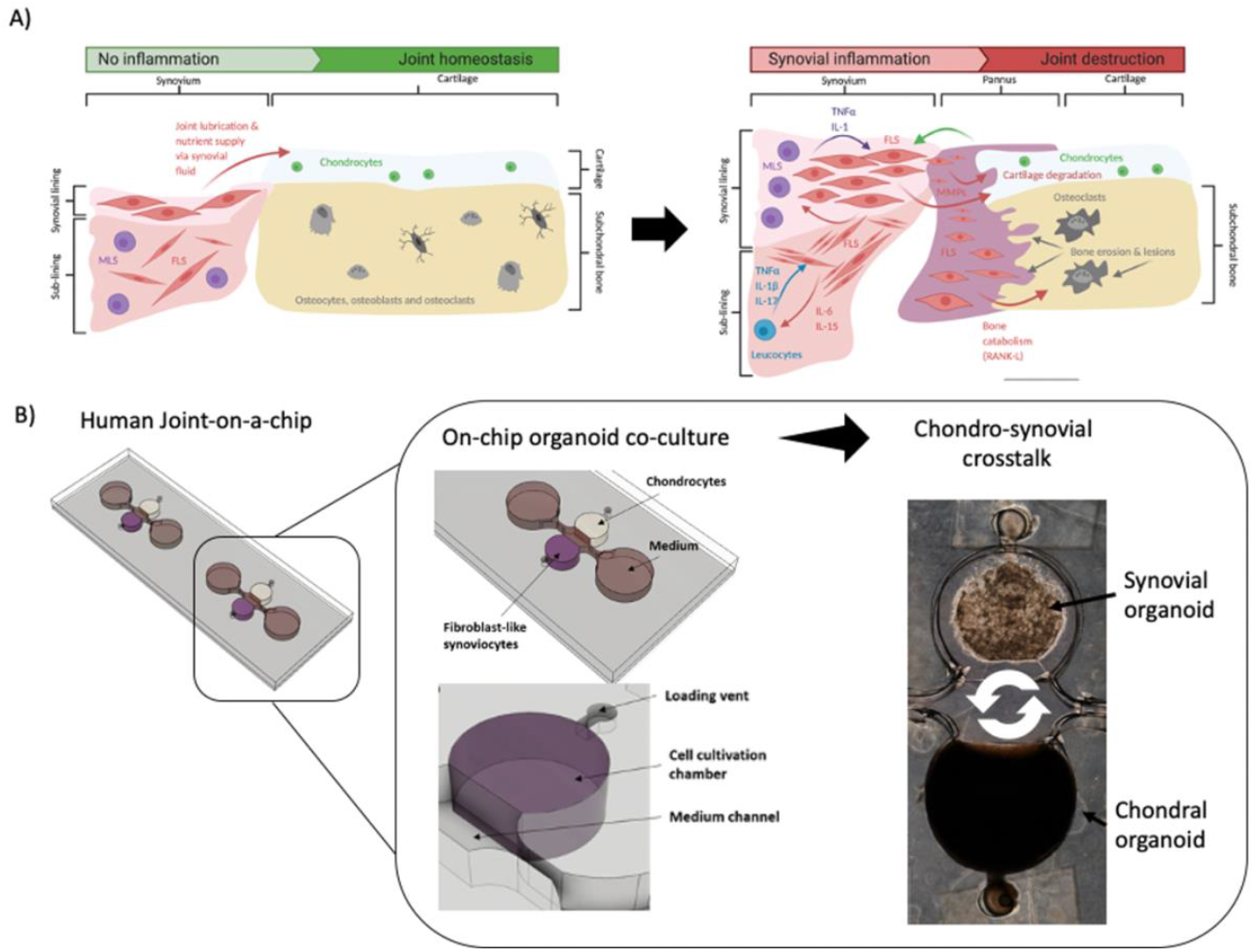
A) Schematic drawing of the synovial activity influencing cartilage and subchondral bone in healthy as well as rheumatoid joints. B) Overview of the organoid-based joint-on-a-chip co-culture system comprising of chondral and synovial compartments for reciprocal inflammatory cross-talk.

Over the last decade arthritis research has gradually moved from conventional two-dimensional to more complex three-dimensional cell cultures, which allowed more physiologically relevant interactions mediated by unidirectional as well as reciprocal cell communication and transformation to human arthritic synovial tissue *in vitro*.^18^ Starting in 2006, Kiener *et al.* introduced a 3D synovial micromass model with FLS embedded in Matrigel to spontaneously form a synovial construct comprising of lining and sub-lining layers.^5^ Broeren *et al.* adapted this Matrigel-based primary model by combining FLS and CD14+ monocytes. When stimulated with TNFα, these micromasses triggered arthritic hallmarks including synovial lining thickening, pro-inflammatory macrophage phenotype (M1) as well as upregulation of pro-inflammatory cytokines.^19^ Maracle *et al.* used a 3D drug screening model based on spheroids fabricated from RA-FLS and endothelial cells to recapitulate synovial angiogenesis, which constitutes a detrimental process in RA disease progression. ^20^ Furthermore, Peck *et al.* successfully engineered a scaffold-free tri-culture drug screening model comprising of FLS (SW982 cell line), LPS-activated macrophages and primary isolated chondrocytes from knee-cartilage to investigate cartilage deconstruction and RA reversal in affected porcine joints.^21^ Our group used a lab-on-a-chip with non-invasive light scattering sensing strategy to rapidly investigate TNFα induced rheumatoid architectural changes in 3D synovial micromass cultures concurring with synovial reorganisation as well as interleukin (e.g. IL-6, IL-8) and MMP (e.g. MMP-1, MMP-3, MMP-13, etc.) molecule secretion.^22^ More recently, also microphysiological models and organ-on-a-chip devices have been introduced to model various aspects of arthritic diseases: Ma *et al.* established a co-culture of human FLS (synovial sarcoma SW982 cell line) with murine pre-osteoclasts (RAW264.7) and bone-marrow-derived stem cells (BMSCs) to study FLS migration and invasion-mediated bone erosion in RA.^23^ Nonetheless, to date most systems rely on non-human and/or immortalised cell lines with no microsystem using primary human cell types to establish these complex 3D co-cultures to model arthritic diseases.

Therefore, herein we present a joint-on-a-chip model based on co-cultivation of two spatially separated hydrogel-based 3D organoid constructs to gain a deeper understanding of how reciprocal tissue cross-talk from synovial to chondral organoids and vice versa stimulates joint (patho)physiology (see Fig. 1B). An essential aspect of our joint-on-a-chip development capable of mimicking the human chondro-synovial niche was concerned with improved reproducibility, which is key to all advanced *in vitro* cell-based assays. Consequently, a range of parameters known to influence organoid formation is investigated, including passaging of isolated synovial cells in 2D, variations of immune cell composition and cytokine secretion. Another focus is the investigation of the effects of chondro-synovial cross-talk on tissue-level behaviour. As a first practical application of our dual organoid culture chip, the addition of chondrogenic growth factor (TGFß3) and fetal calf serum (1% or 10% FCS) was investigated to assess changes in pro-inflammatory cytokine secretion. Overall, the proposed chip-based organoid co-culture model shows promising results due to a high degree of microenvironmental and cellular control and excellent imaging capabilities of the architectural context and cellular composition not feasible for conventional 3D micromass and spheroid cultures established with microtiter plates.

## Experimental

### Clinical Specimens

Human synovial tissues from patients were obtained from RA undergoing synovectomy (fulfilling the ACR/EULAR classification criteria^24^) and an OA patient with written informed consent. Under the terms of the Medical University’s ethics committee in Vienna (EK-No. 1065/2011). The medical background of patients was documented and included age (average age: 55, min.: 27, max.: 71), sex (8 female and 2 male), tissue location (carpals: 4, phalanges: 3, elbow joint: 2, tibiofemoral joint: 1), diagnosis (total: 10, rheumatoid arthritis: 9 patients, osteoarthritis: 1 patient) and medication (no treatment: 2, Embrel: 2, Arava: 1, cortisone: 3, Salazopyrin: 1, Arava + cortisone: 1). Primary cell populations were isolated and stored in liquid nitrogen, as described earlier.^25^

### Cell Culture

FLS were thawed at 37 °C and immediately plated in a T75 cell culture flask containing pre-warmed DMEM culture medium supplemented with, 10% FCS, 1% MEM NEAA and 1% antibiotics. Cells were split upon reaching 90% confluency. Before passaging, 1 ml of culture medium was collected and frozen for supernatant analysis. For passaging experiments, cells were enzymatically detached, pelleted and resuspended in culture medium. An aliquot of 5*10^5^ cells was weekly transferred to a new flask, whereas the residual cells were used for flow cytometry analysis and 3D cultivation.

Commercial human primary chondrocytes (C-12710; female donor; PromoCell) were maintained in chondrocyte growth medium (411-500; CellApplications Inc.) and was exchanged every other day.

### Synovial micromass culture in well plates

Well plates were coated with 0.03 w/v% ethanolic poly(2-hydroxyethyl methacrylate (poly-Hema) as described initially for synovial micromass cultures.^5^ Briefly, after pelleting the primary FLS cells at a passage number between 1 and 8 were resuspended in ice-cold Matrigel at a concentration of 3000 cells/μl Matrigel. To initiate micromass cultures, a 20 μl drop of the cell-laden Matrigel suspension was pipetted into the centre of a coated well. After 40 minutes of polymerisation, cell/Matrigel drops were overlaid with 2 ml fully supplemented organoid culture medium containing 1% Insulin-Transferrin-Selenium (ITS) and 0.032 μg/ml L-Ascorbic acid 2-phosphate with a weekly medium exchange.

### Microfabrication

The two multi-layered designs for either synovial monoculture or chondro-synovial co-culture illustrated in ESI Fig. 1 were fabricated using a CAMM-1 GS-24 (Roland, Germany). They assembled layer by layer by xurography from two-dimensional CAD templates from 500 μm thick transparent silicone sheets (MVQ Silicone) as described previously.^26,27^ The basic design principles were adapted from an earlier vascular organoid study^28^, and circular medium reservoirs were added to contain a total medium volume of 300 μl. For height variation, the layer numbers of the circular hydrogel compartment were adjusted in the range of 1-4 mm. To prevent cellular outgrowth from within organoids, chip surfaces were coated with 0.5% ethanolic Lipidure^®^ (amsbio, UK) overnight before UV light sterilisation for 1 hour.

Two basic designs with five structured PDMS layers were assembled on object slide base layers to form a joint-on-a-chip (see ESI Fig. 1A): The monoculture design (left column layers I-V) comprised of a single organoid chamber (Ø 5 mm), whereas two opposing organoid chambers allow for soluble tissue-level communication via reciprocal signalling (right column).^28,29^ To improve medium replacement and sample withdrawal of the optimised design, the medium supply channels interconnect two circular medium reservoirs. To enable reproducible and facile loading of hydrogels structured layers I and II form a step-like hydrogel stopper feature that prevents hydrogel bleeding into the medium supply channel. Additionally, structure layers III-V form a chamber geometry that favours controlled organoid formation by an anchoring organoid condensation guide. (see ESI Fig 1BC for detailed functional and dimensional elaborations on both features).

### On-chip organoid culture

For the generation of synovial organoids, an FLS-containing hydrogel suspension between passage number 1 and 8 was loaded via loading vent into each cell cultivation chamber. To start organoid formation, a total of 45 μl FLS cells in Matrigel was slowly injected at a concentration of 3000 cells/μl through the hydrogel port and was sealed by a small piece of PCR sealing film on ice. After 40 minutes polymerisation at 37 °C, organoid chip medium was added to start the organoid culture. To establish on-chip co-cultures, a previously established cartilage organoid based on TISSEEL Fibrin hydrogel (Baxter, Austria) was injected in the chondral compartment as previously described.^29^ Briefly, 30 μl of Fibrinogen solution at a concentration of 100 mg/ml containing 2000 cells/μl was mixed with 30 μl 4U/ml thrombin solution supplemented with 40 μM CaCl_2_ solution and was polymerised for 30 min at 37 °C. For medium composition experiments, TGFβ3 was added to the chip medium at 1 and 10 ng/ml in either 1% or 10% FCS-containing organoid culture medium and compared to commercial chondrogenic differentiation medium (411D-250; CELL Applications Inc., USA).

### Flow cytometry

Two different staining mixes were prepared on ice, as shown in ESI Table 1. Aliquots of 1.5*10^5^ cells were washed with flow cytometry buffer (PBS with 2% FCS) and pelleted at four °C and 1350 rpm for 8 min. After pelleting, cells were resuspended in 50 μl of the staining mix solution, incubated for 20 min, washed and resuspended in 150 μl flow cytometry buffer. Measurements were performed on a BD Canto II flow cytometer using *BD FACS Verse™.* (Note: One measurement cycle lasted either 100 000 cells or 2 minutes.) Gating of cell populations was performed as described elsewhere.^30,31^

### Confocal microscopy

Imaging was performed using an upright Leica TSC SP5 Microscope (Leica Microsystems) with 20x and 40x water immersion objectives. Imaris software (Bitplane) was used for image processing. Live-cell staining was performed with fluorescent cell tracker dyes (Green CMFDA/Orange CMRA and Deep Red; Thermo Fisher) before pelleting and hydrogel mixing according to the manufacturer’s staining protocol and ESI Table 2.

### Morphometric analysis

For morphometric analysis, manual tracing of the organoid boundaries of phase-contrast images of organoids taken at selected time points was used in ImageJ (Version 2.0.0.rc.69/1.52.p) in combination with the shape descriptors option of the ‘measure’ function as previously described.^32^

### Supernatant protein analysis

For AlphaLISA analysis of IL-6, IL-8, MMP-1 and MMP-3 (AL223C, AL224C, AL284C, AL242C, Perkin Elmer) undiluted supernatants of 2D cultured FLS were analysed according to the manufacturer’s protocol using an EnSpire^®^ Multimode plate reader (Perkin Elmer).

For cytokine analysis of IL-6, IL-8, MMP-13 and VEGF using human Magnetic Luminex Assay (R&D Systems), supernatants of synovial and chondral organoids in mono and co-culture were diluted 1:10 in dilution reagent and analysed according to the manufacturer’s protocol using a Luminex™ MAGPIX™ Instrument System (Invitrogen). After medium background subtraction, molecule secretion of mono- and co-cultures were normalised to 10^4^ cells per mL and day to compensate for differences in total cell numbers between synovial and chondral monocultures as well as co-cultures.

## Results & discussion

### Characterisation of the chip-based three-dimensional synovial organoid model

Initially, individual hydrogel compartments were studied separately to individually investigate the formation and maturing of each organoid unit. In case of the synovial organoid cultures, RA patient derived primary FLS are mixed into Matrigel prior injection and cultured up to four weeks. It is important to note that during synovial organoid formation, significant hydrogel condensation and remodelling takes place. For instance, Fig. 2AB shows the influence of increasing hydrogel volume (Note: Increasing chamber heights from 1 to 4 mm results in a volume increase from 19.6 to 78.5 μl.) on condensation behaviour, where RA-FLS organoid volume gradually declined between 18-36% after 28 days of organoid cultivation. Interestingly, a chamber height of 1 mm (19.6 μL) was insufficient to generate a single synovial organoid; instead fragmentation of the organoids and cell dispersion occurred; see also images of ESI Fig. 2. In contrast, chamber heights of 1.5, 2.0 and 2.5 mm displayed uniform organoid formation exhibiting an area of 3.8 ± 0.27 mm^2^ at day 28. Larger volumes (e.g. chamber heights of 3 and 4 mm) produced bigger synovial constructs with 5.7 ± 0.12 mm^2^ and 7.1 ± 0.09 mm^2^, respectively. A more detailed morphology analysis revealed that most homogenous synovial organoid formation is achieved using a cultivation chamber with 2 mm height, where almost spherical structures are obtained. Fig. 2C illustrates the time-dependent change of synovial organoid shape from rectangular to spherical throughout 28 days of cultivation period. In a follow-on control study, on-chip synovial condensation dynamics is compared to polyHEMA-coated anti-adhesion well plates (MMC). Results of this comparative study are shown in Fig. 2D und 2E and reveal similar architectural features with a significant higher reproducibility exhibiting a 3.5-fold lower relative standard deviation of 4.4% (n=5) compared to 15.5% (n=3) for MMC cultures generated in anti-adhesion well plates.

**Fig. 2.**
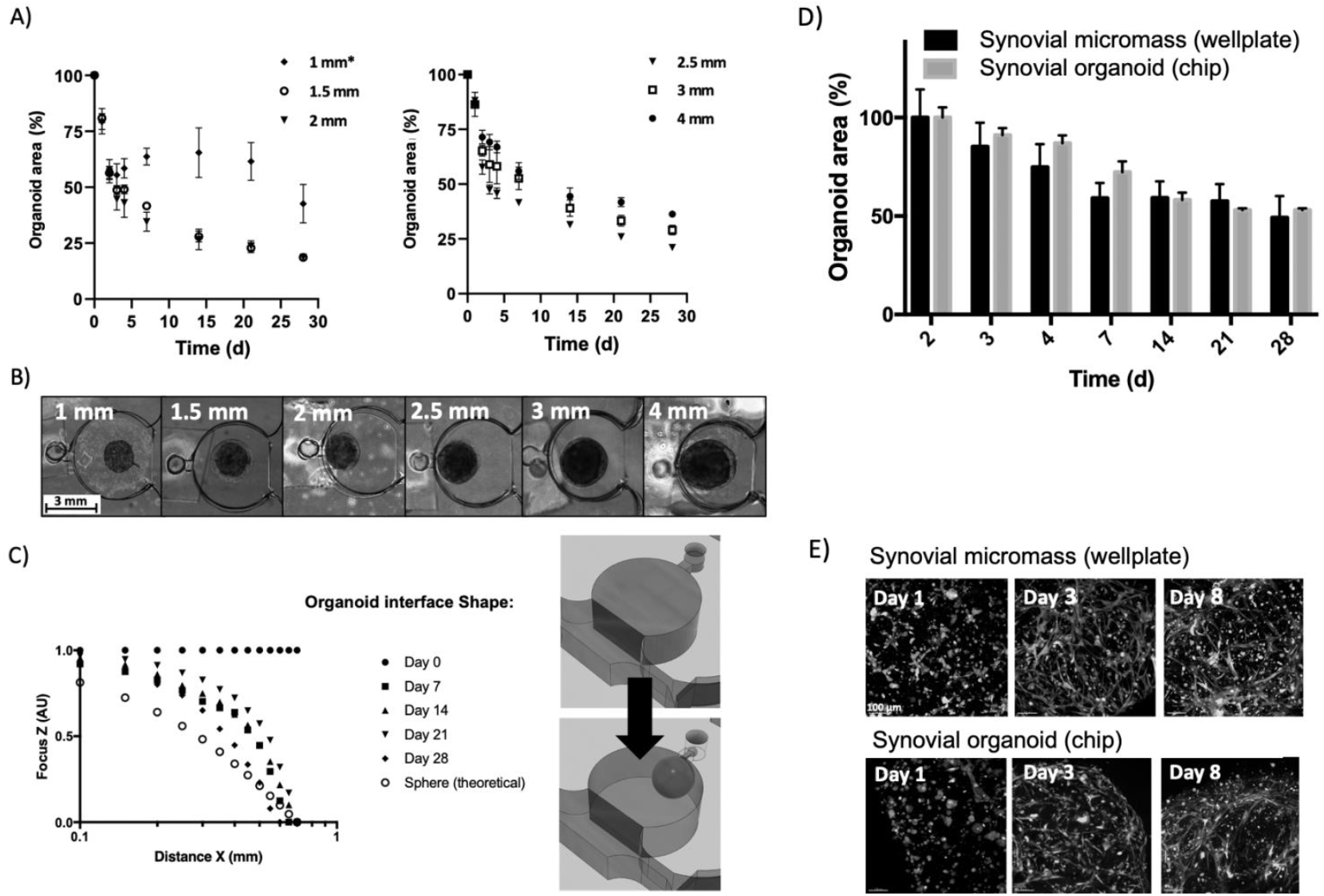
Characterization of the formation of synovial on-chip organoids with RA patient-derived fibroblast-like synoviocytes. A) Influence of organoid chamber height on organoid area throughout 28 days of cultivation. (Data is expressed as mean ± standard deviation for n=3 organoids of a single donor RA synoviocyte sample.) B) Representative brightfield images of synovial organoids formed from various chamber heights at day 28 post-seeding. C) Change of synovial organoid shape from rectangular to spherical throughout 28 days of cultivation in 2 mm high organoid compartments using manual z focus analysis of the organoid surface. D, E) Comparison of organoid area over 28 days and initial synovial network formation up to day 8 for on-chip organoids and wellplate micromass cultures. (Data is expressed as mean ± standard deviation for n=3 organoids generated from a single RA synoviocyte donor sample.)

### Influence of passaging and cultivation on the heterogeneity of patient-specific synovial cell populations

Arthritic synovial tissue comprises aside from synovial fibroblasts (FLS) a diverse set of immune cells such as macrophages, B cells and T cells. While co-cultivation of primary synovial fibroblasts and immune cell populations under standard 2D culture conditions is known to rapidly deplete immune cells resulting in pure synovial fibroblasts (FLS),^19^ less is known about the effect of this immune cell depletion on the formation of FLS organoids. To evaluate the long-term effect of co-cultivating isolated patient derived FLS and immune cells on synovial tissue heterogeneity, eight individual isolated patient samples (e.g. 7 RA and 1 OA patient) were characterised using flow cytometry and supernatant analysis. Results of this study are shown in ESI Fig. 3AB, indicating a gradual depletion of leucocytes and selective growth of synovial fibroblast over 5 weeks (5 passages). A high degree of interpatient immune cell variability was evident for CD45^+^ leucocytes ranging from 22% to 1.1% between individual patient samples. Further analysis showed variations of sub-population of 4.1% to 0.1% CD3^+^/CD14^-^ T cells, and 3.9% to 0.01% CD19^+^/CD14^-^ B cells as well as a predominant population of 8.9% to 0.6% CD14^+^/CD3^-^ macrophages. Similarly, a gradual decrease of pro-inflammatory cytokines (e.g. IL-6 and IL-8) and degradative matrix metalloproteinases (e.g. MMP-1 and MMP-3) in cell culture supernatant was detected over the five-week cultivation period (see ESI Fig. 3C).

Next, the impact of medium composition for on-chip organoid cultivation and 2D/3D cultivation on immune cell survival was investigated in more detail. Results shown in Fig. 3A revealed a higher degree of leucocyte survival when substituting the DMEM culture medium with leucocyte medium (e.g. RPMI 1640) leading to 4.9, 2.4 and 1.8-fold higher T cell, macrophage and B cell subpopulations after two weeks of sub-cultivation. In turn, synovial organoid on-chip cultivation further improved leucocyte survival for both medium types (n=2; see Fig. 3B) with 11- and a 20-fold increase in leucocyte populations compared to conventional 2D cultivation, that showed a declining population trend. First of all, this means that both the amount and type of immune cells present in rheumatoid arthritis patients vary significantly and need to be accounted for in an *in vitro* arthritis model. Propagated as 2D culture in DMEM medium, immune cell depletion decreases the arthritic sample heterogeneity regarding cell composition and cytokine secretion and favours FLS growth, whereas p0-1 synovial populations reflect an arthritic and patient-specific environment with varying cell populations of the immune cell infiltrate, thus also variations in the respective cytokine secretion profile. Furthermore, our chip-based synovial organoid system featuring a three-dimensional tissue-like microenvironment protects synovial leucocyte populations from depletion even in unfavourable culture media (e.g. DMEM).

**Fig. 3.**
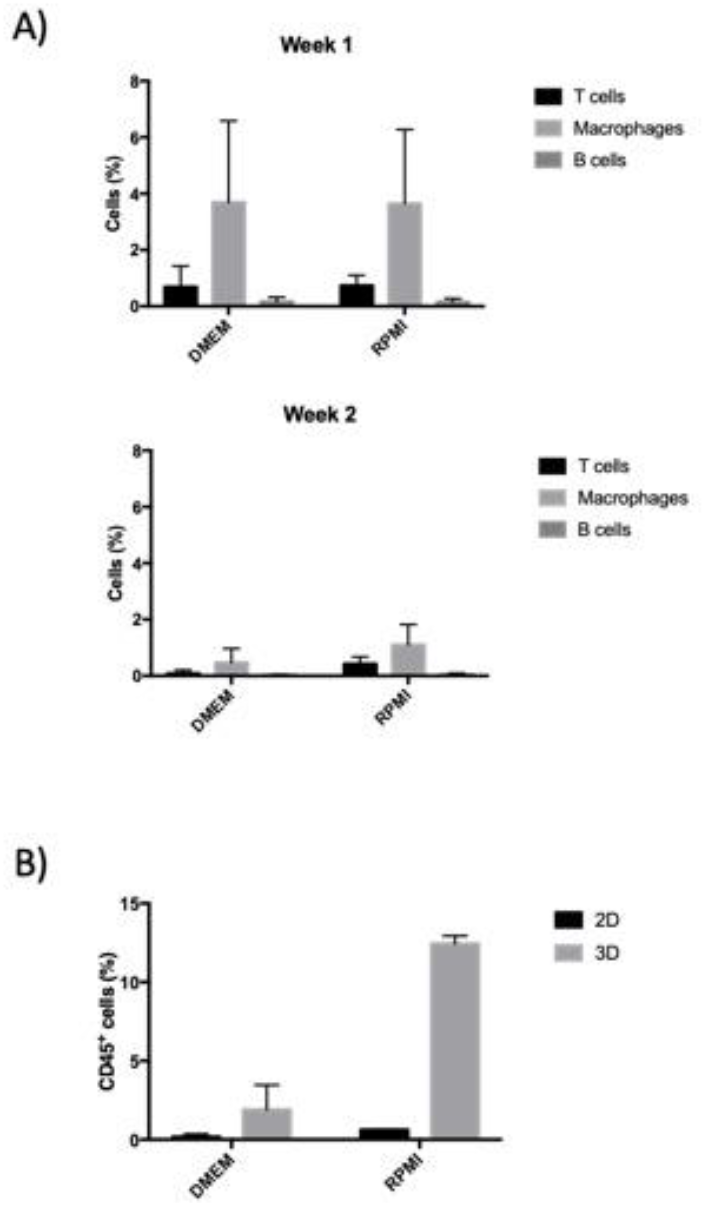
A) Impact of medium type on leucocyte populations within synovial RA-FLS organoids. (n=2; data is expressed as mean ± standard deviation in % of total CD 45 positive cell populations). B) Comparison of CD45 positive leucocyte populations from isolated RA synovia in p0 cultured as 2D cultures or 3D organoids in DMEM and RPMI 1640 medium using flow cytometry analysis. (n=2; data is expressed as mean ± standard deviation in %).

**Fig. 3.**
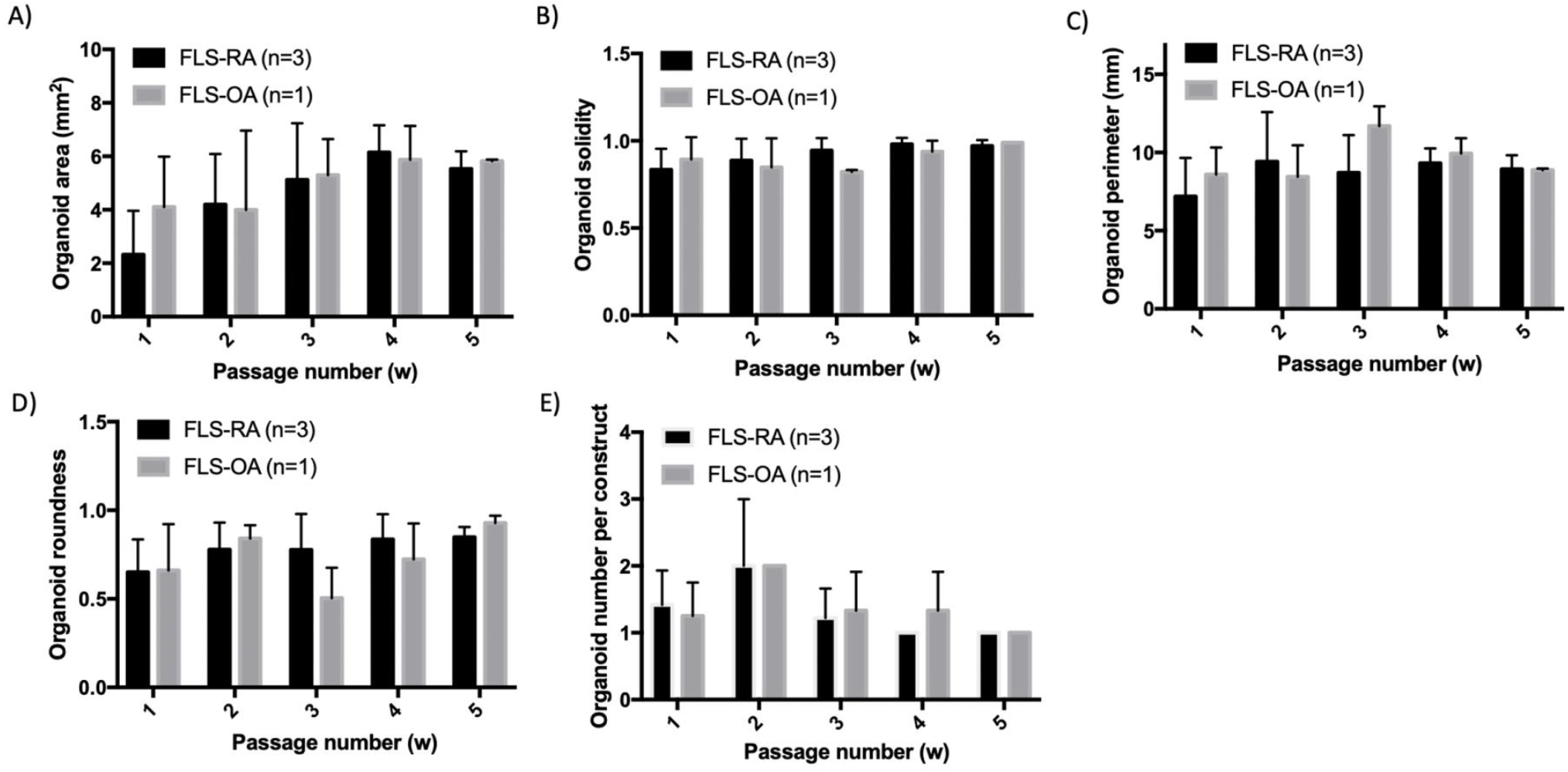
Morphometric analysis of synovial organoids produced with fibroblast-like synoviocytes from increasing passage numbers after 28 days of cultivation. (A-D) Influence of fibroblast-like synoviocyte passage number and cell origin on morphology parameters including A) area, B) solidity, C) perimeter and D) roundness. E) Impact of passaging on organoid number per compartment. (RA; Rheumatoid arthritis. OA; osteoarthritis. Data is expressed as mean ± standard deviation for n=3 organoids generated from three individual RA and a single OA patient donor samples.)

### Impact of passaging on the formation of chip-based synovial organoids

To further understand the effects of synovial cell passaging on chip-based 3D synovial cultures, organoid formation and secretion of cytokines were investigated in ensuing experiments. On-chip generated synovial organoids of increasing passages (p1-5) were cultivated over five weeks, and vital organoid parameters including area, solidity, perimeter, roundness and organoid numbers per unit were analysed. Fig. 4 and ESI Fig. 4 show morphometric analyses at day 28 post-seeding and point at stable organoid formation peaking at 5.5±0.6 mm^2^ and 5.8±0.06 mm^2^ for later passages in contrast to freshly isolated FLS populations. Interestingly a high inter-sample variability within individual RA (n=3) and OA (n=1) patient-derived synovial organoids were observable. Early passages (p1) showed a tendency to form smaller cell aggregates ranging from 1.8 - 4.1 mm^2^ with higher variability (0.42% RSD). In turn, FLS organoids formed from p5 cells yielded reproducible organoid areas of 5.6±0.67 mm^2^ (0.01% RSD; see Fig. 4B). Remaining morphometric parameters, including perimeter (0.04% RSD), roundness (0.05% RSD) and solidity (0.01% RSD) further pointed at highest reproducibility in the presence of higher passage numbers. A direct comparison between RA and OA-derived organoids (for details see ESI Table 3) revealed homogenous organoid shapes independent of patient or disease origin. However, morphometric scores (e.g. solidity and roundness) were slightly better for the OA-derived FLS after five passages compared to the RA organoids that already showed good formation performance for earlier passages (p3-4). A similar trend was obtained for the system’s reproducibility to form a single organoid in each individual organoid compartment (see Fig. 4E), which scored best for both OA and RA samples after 5 weeks of passaging. Overall, these results suggest that the purity of FLS populations is improving organoid formation reproducibility due to immune cell depletion.

**Fig. 4.**
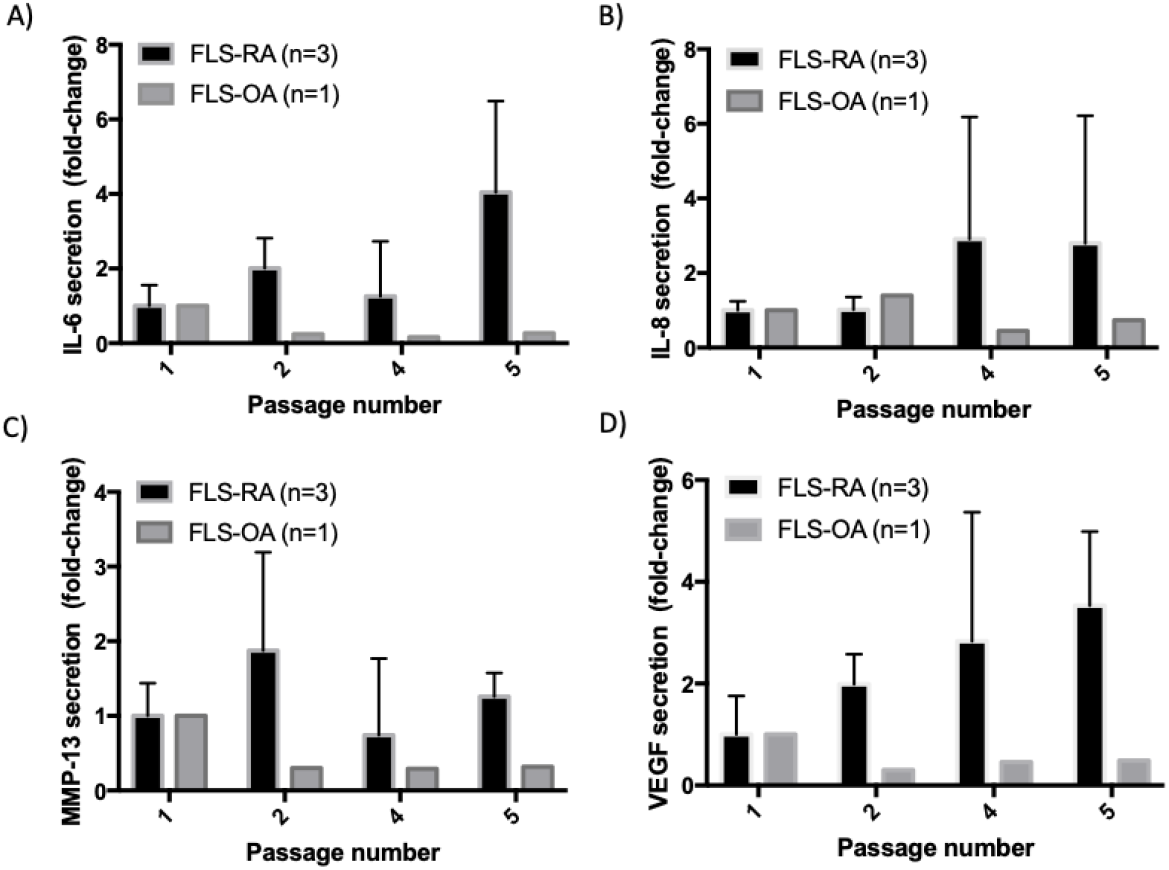
Secretion analysis of synovial organoids at day 28 post-seeding produced with fibroblast-like synoviocytes from increasing passage numbers for A) IL-6, B) IL-8, C) MMP-13, and D) VEGF using Luminex multiplex technology. (RA; Rheumatoid arthritis. OA; osteoarthritis. Data is expressed as mean ± standard deviation for n=1-2 organoids generated from three individual RA and a single OA patient donor sample.

Since on-chip cultivation of artificial tissues is known for accumulation and retention of secreted molecules^33,34^, secretion analysis from collected supernatants of the above organoid cultures were conducted to monitor secretory behaviour for FLS of increasing passages. Results from our comparative secretion analysis of IL-6, IL-8, MMP-13 and VEGF up to passage p5 are shown in Fig. 5A-D. While OA-derived organoids showed a decreasing secretion trend for increasing passages, RA-derived organoids gradually increased molecule secretion over time when cultivated in a three-dimensional organoid system. For instance, RA-FLS organoids showed 2.7-, 4, 1.25 and 3.5-fold higher secretion at day 28 for IL-6, IL-8, MMP-13 and VEGF for p5 compared to p1 RA-FLS cells. In contrast, OA-FLS decreased their secretion levels by 26%, 73%, 69% and 52% for IL-6, IL-8, MMP-13 and VEGF at p5, respectively. Even though the relative secretory pattern over 5 passages was similar, when analysing organoid supernatants collected at day 7 and 28 post-seeding the absolute secretory values were higher in the first week of organoid culture, where cell populations are still reorganizing organoid architecture as synovial networks (see ESI Fig. 5) and declined in the following two weeks.

**Fig. 5.**
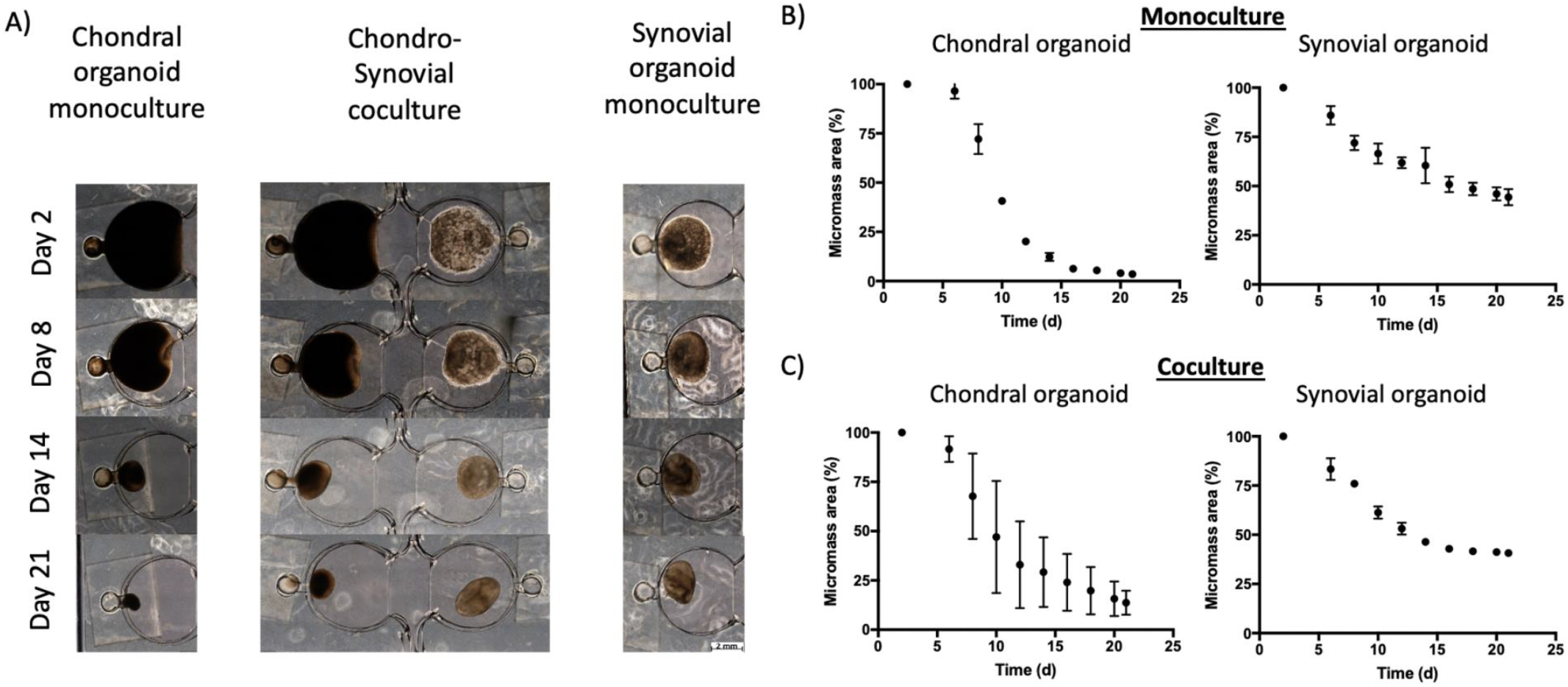
Impact of organoid co-culture on organoid formation dynamics over three weeks of co-cultivation. Representative images A), as well as ImageJ organoid area analysis of B) on-chip monocultures in comparison to C) chondro-synovial organoid co-cultures. (Scale bar = 2mm. Data is expressed as mean ± standard deviation for n=2 organoids from individual RA synoviocyte and ‘healthy’ chondrocyte donor samples.)

Overall, due to the fact that organoids from older passages showed stable organoid formation while retaining similar molecule secretion profile, patient-derived RA-FLS cells from p5 on were used in subsequent experiments to combine synovial organoids with a chondral unit inside our chip-based platform for tissue-level cross talk.

### Establishment of a chondro-synovial joint-on-chip organoid co-culture model

Another critical aspect in establishing a functional *in vitro* arthritis model of the joint is concerned with proper tissue to tissue interactions between the synovium as inflammatory tissue and the cartilage as target tissue during the onset and progression of the disease. Next, co-cultures of synovial (patient-derived RA-FLS cultures) and chondral (primary human chondrocytes) organoids were established and compared to on-chip cultures of individual organoid units (see Fig. 6A). It is important to note that chips containing only chondral organoids (chondrocytes suspended in 50 mg/ml Fibrin) showed typical round chondrocyte morphology within the fibrin construct (see ESI Fig. 6). Chondral organoid monocultures condensed to 3.5±1.9% of the initial organoid area, whereas synovial monocultures retained 44.4±4.1% organoid area after 28 days of culture (see Fig. 6B). In co-culture, synovial organoids condensed slightly more down to 40.7±0.3% organoid area. In contrast, the chondral compartment was affected more by tissue cross-talk showing 4-fold less condensation, resulting in 13.7±4.1% organoid area. This means that both organoids were affected by co-cultivation: Chondral organoids showed increasing and synovial model decreasing organoid area compared to their respective monocultures. Additionally, organoid co-culture systems yielded already elevated cytokine secretion (IL6 and IL-8; see Fig. 7A-D) at day three that gradually decreased over time, while other inflammatory responses such as the MMP-13 response increased by 2-to 3-fold throughout three weeks of co-cultivation. Interestingly, chondral organoids in co-culture showed a decrease in VEGF secretion compared to chondral monocultures pointing at a more physiological chondral environment^35,36^, which may explain the increase in organoid area.

**Fig. 6.**
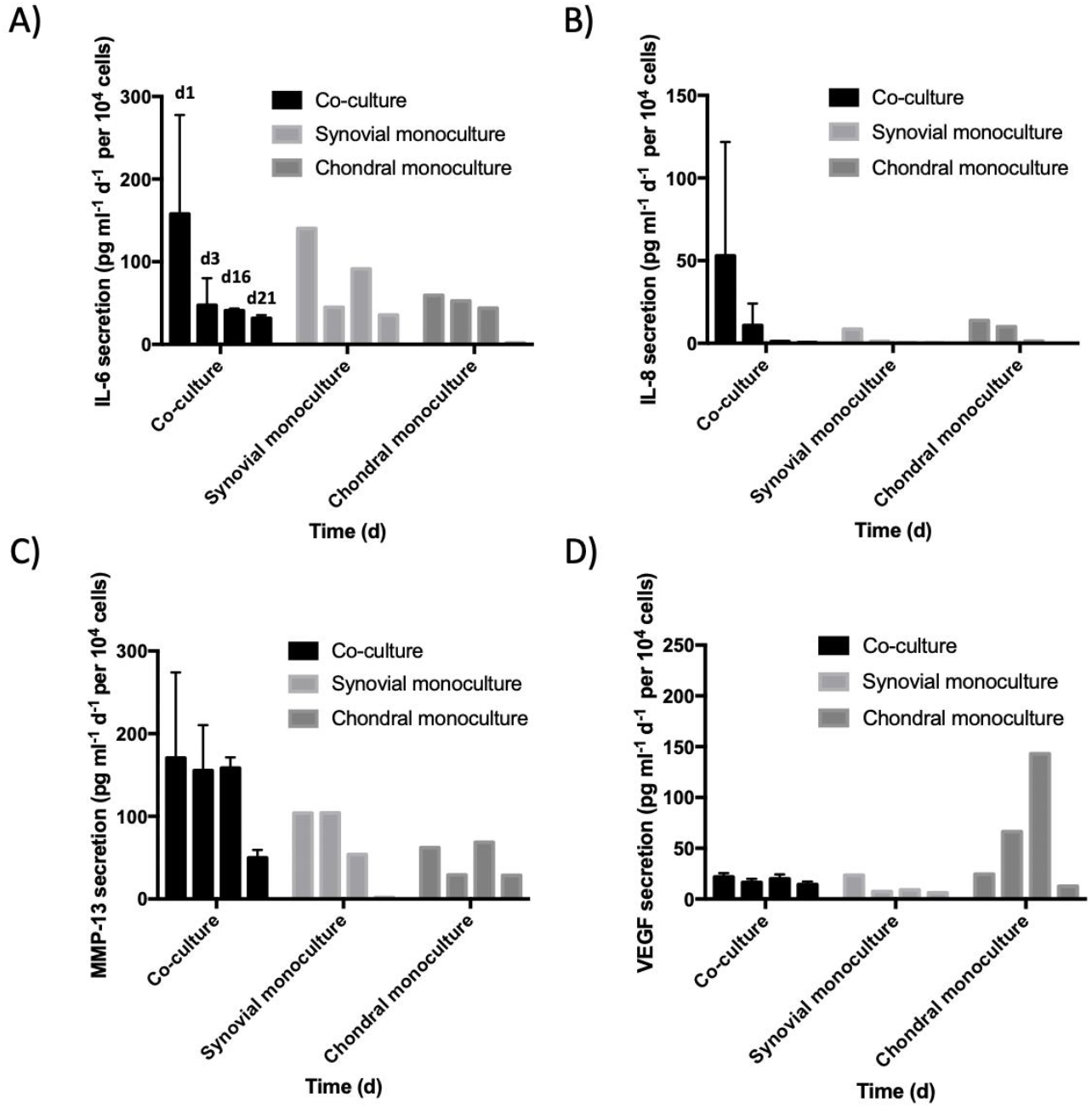
Impact of co-culture on molecule secretion for on-chip chondro-synovial organoid co-cultures compared to organoid monocultures of RA patient origin on at day 21 post-seeding for A) IL-6, B) IL-8, C) MMP-13, and D) VEGF at day 3, 6, 16 and 21 post-seeding. (Normalized data is expressed as mean ± standard deviation for n=1-2 for organoid co-cultures generated from individual synoviocyte and chondrocyte donor samples.)

**Fig. 7.**
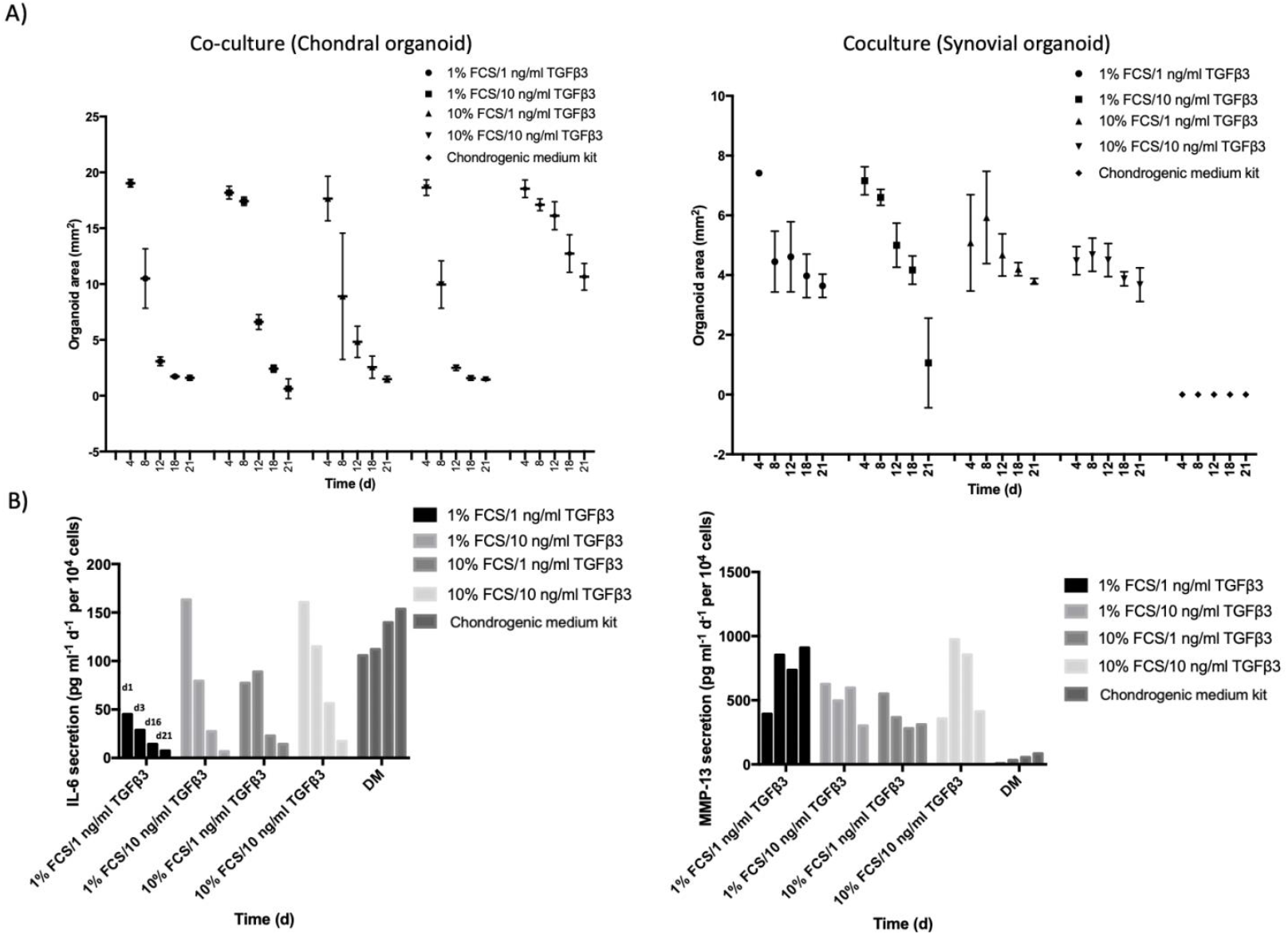
Impact of medium type and supplementation on RA-organoid formation and secretion profile of chondro-synovial organoid cultures. A) Progression of organoid area of chondro-synovial organoid co-cultures (n=3-4), and B) secretion analysis of IL-6 and MMP-13 (n=1) of co-cultured chondro-synovial organoids for TGF-ß3 supplemented DMEM as well as chondrogenic differentiation medium at day 3, 6, 16 and 21 post-seeding using Luminex multiplex analysis. (Data is expressed as mean ± standard for organoid co-cultures generated from individual synoviocyte and chondrocyte donor samples.)

To demonstrate that the current chondro-synovial chip system can be used for optimisation and screening efforts, the effect of changing medium compositions such as 1% vs. 10% FCS and TGFß3 supplementation (e.g. 1 ng/ml vs. 10 ng/ml) on organoid formation was investigated in subsequent experiments. Results in Fig. 8A and ESI Fig. 7 show stable chondral organoid formation retaining up to 95±2% of the initial organoid size in the presence of chondrogenic differentiation medium. The presence of 1% FCS combined with 10 ng/ml TGFß3 exhibited similar values at day 8 than the commercial chondrogenic medium that retained 92±3%. All other growth media compositions showed faster chondral condensation resulting in 50-55% of the initial organoid area. Interestingly, while the chondral organoids showed a lesser degree of condensations when cultivated with chondrogenic differentiation medium, the synovial organoid formation was halted. The combination of 1% FCS and 10 ng/ml TGFß3 introduced overall progressive and strongest tissue condensation down to 14.8±20.9% in both organoids. It has to be noted that even though TGFß3 is frequently used for chondrogenic maintenance of primary chondrocytes, it has been demonstrated to induce synovial fibrosis *in vitro*.^19^ Therefore, this increasing organoid instability may be linked to fibrotic molecular events. Final, secretion analysis (see Fig. 8B) revealed that organoid co-cultures responded with increasing IL-6 and low MMP-13 levels for chondrogenic medium, whereas any other medium supplements showed declining IL-6 with high MMP-13 levels. These results demonstrate that the proposed co-culture model is susceptible to changes in medium composition and growth factor stimulation with alterations in architecture and molecular patterns based on reciprocal tissue-level cross-talk. Consequently, the presented chip-based technology paves the way to create and use more complex and holistic 3D models in arthritis research.

## Conclusions

In the current study, we have for the first time to our knowledge established a chip-based chondro-synovial dual organoid model based on self-organising patient-derived cells that allow investigating reciprocal tissue-tissue cross-talk in orthopaedic/arthritic research. Due to the intrinsic microfluidic design features, the current microfluidic joint-on-a-chip design enables the reliable formation of organoids with high reproducibility and positioning precision, which is a crucial aspect for any organ-on-a-chip analysis system used in biomedical and biopharmaceutical research. We also demonstrated that expanded and purified synovial fibroblast cultures generate organoids of higher precision than freshly isolated cell populations due to immune cell depletion and loss of inflammatory secretome. Moreover, we showed that co-cultivation of chondral and synovial organoids improved chondral organoid stability compared to individual organoid monocultures. As a first practical application of our joint-on-a-chip systems differential and dynamic morphological and secretory changes were shown when challenged with different cultivation environments and inflammatory factors either favouring chondral or synovial organoids. We believe that our current dual organoid-based joint-on-a-chip approach highlights the importance of tissue-level cross-talk in the context of musculoskeletal diseases, including RA and OA.

## Supporting information

Supplementary Information: ESI Table 1-3, ESI Fig. 1-7

## Author contributions

AF, EIR, IOC, MR, RAB, and SS performed the experiments and analysed the data. F. S. and J. H. provided primary patient tissue. SS, BB, WH and HR provided expertise and assistance with fibrin-based organoids. H. P. K., MR and P. E. supervised the work. MR, PE, HPK, RW and ST edited the manuscript.

## Acknowledgements

This project was supported by the Austrian Federal Ministry of Education, Science and Research (BMBWF; Grant agreement number 1612889) and the Vienna Science and Technology Fund (WWTF, project number LS13-092).

## Notes and references

^†^These authors contributed equally.

