## Supplementary Information: ESI Table 1-3, ESI Fig. 1-7 for "Establishment of a human three-dimensional chip-based chondro-synovial co-culture joint model for reciprocal cross-talk studies in arthritis research"

[a.] *Karl Chiari Lab for Orthopaedic Biology (KCLOB), Department of Orthopedics and Trauma Surgery, Medical University of Vienna, Währinger Gürtel 18—20, 1090 Vienna, Austria*
[b.] *Faculty of Technical Chemistry, Vienna University of Technology, Getreidemarkt 9, 1060 Vienna, Austria.*
[c.] *Division of Rheumatology, Department of Medicine III, Medical University Vienna, Währinger Gürtel 18—20, 1090 Vienna, Austria*
[d.] *Division of Orthopedics, Department of Orthopedics and Trauma Surgery, Medical University of Vienna, Währinger Gürtel 18—20, 1090 Vienna, Austria*
[e.] *AUVA Research Centre, Ludwig Boltzmann Institute for Experimental and Clinical Traumatology, 1200 Vienna, Austria*
[f.] *Austrian Cluster for Tissue Regeneration, 1200 Vienna, Austria*
† These authors contributed equally.

Content: ESI Table 1-3, ESI Fig. 1-7

### Tables

ESI Table 1: Overview over fluorescent staining panels for flow cytometry analysis.

| ***Staining panel 1*** | ***Staining panel 2*** |
|---|---|
| CD90-PerCP-Cy5.5 | CD45-PECy7 |
| CD34-FITC | CD3-APC-Cy7 |
| Podoplanin-PE | CD4-FITC |
| CD45-PECy7 | CD8-PE |
| | CD14-PerCP-Cy5.5 |
| | CD19-APC |

ESI Table 2: Staining and imaging data for different cell tracker dyes from Thermo Fisher.

| ***Cell Tracker™*** | ***Volume (µl/ml)*** | ***incubation time I (min)*** | ***incubation time II (min)*** | ***Ex. (nm)*** | ***Em. (nm)*** |
|---|---|---|---|---|---|
| *Green CMFDA* | 2.5 | 30 | 30 | 492 | 517 |
| *Orange CMRA* | 2.5 | 30 | 30 | 548 | 576 |
| *Deep Red* | 2 | 15 | - | 630 | 660 |

ESI Table 3: Morphometric parameters of synovial organoids triplicates produced from three RA and one OA patient tissue sample.

| ***Tissue origin*** | ***Solidity (±SD)*** | ***Perimeter (in mm ±SD)*** | ***Roundness (±SD)*** |
|---|---|---|---|
| *RA (n=3)* | **0.97±0.03** | **8.93±0.89** | **0.85±0.05** |
| *OA (n=1)* | **0.99±0.005** | **8.87±0.10** | **0.93±0.04** |
| *Best theoretical values* | 1 | N/A | 1 |

### Figures

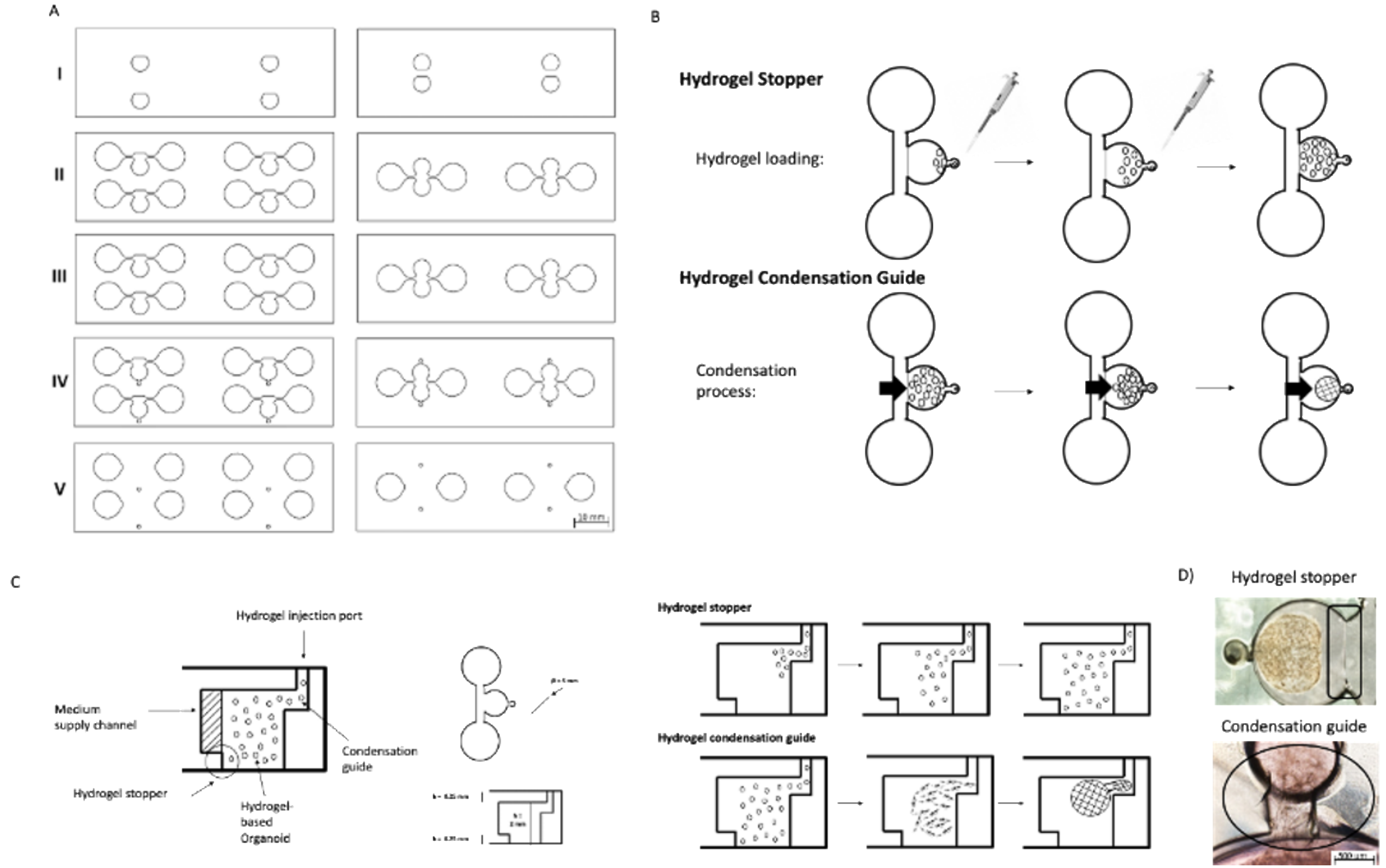


ESI Fig. 1. A) CAD design of the individual structured chip layers from bottom (layer I) to top (layer V). B,C) Top and side-view of the hydrogel stopper and condensation structures within an individual organoid mono-culture unit during seeding (top row) and organoid formation (bottom row) with D) representative microscopic images.

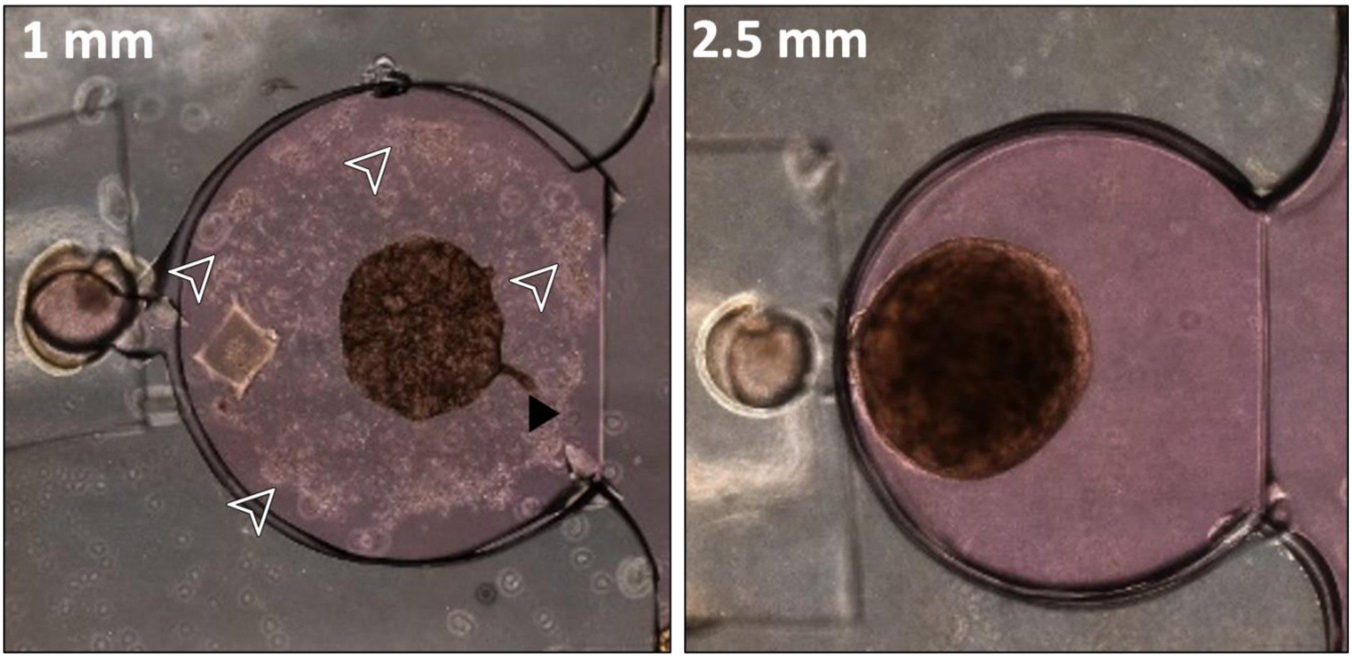

ESI Fig. 2 Fragmentation (black arrowhead) and cell dispersion (white arrowheads) artifacts during FLS organoid maturation in hydrogel chambers of 1mm (left) compared to 2.5 mm (right) height after 28 days post-seeding.

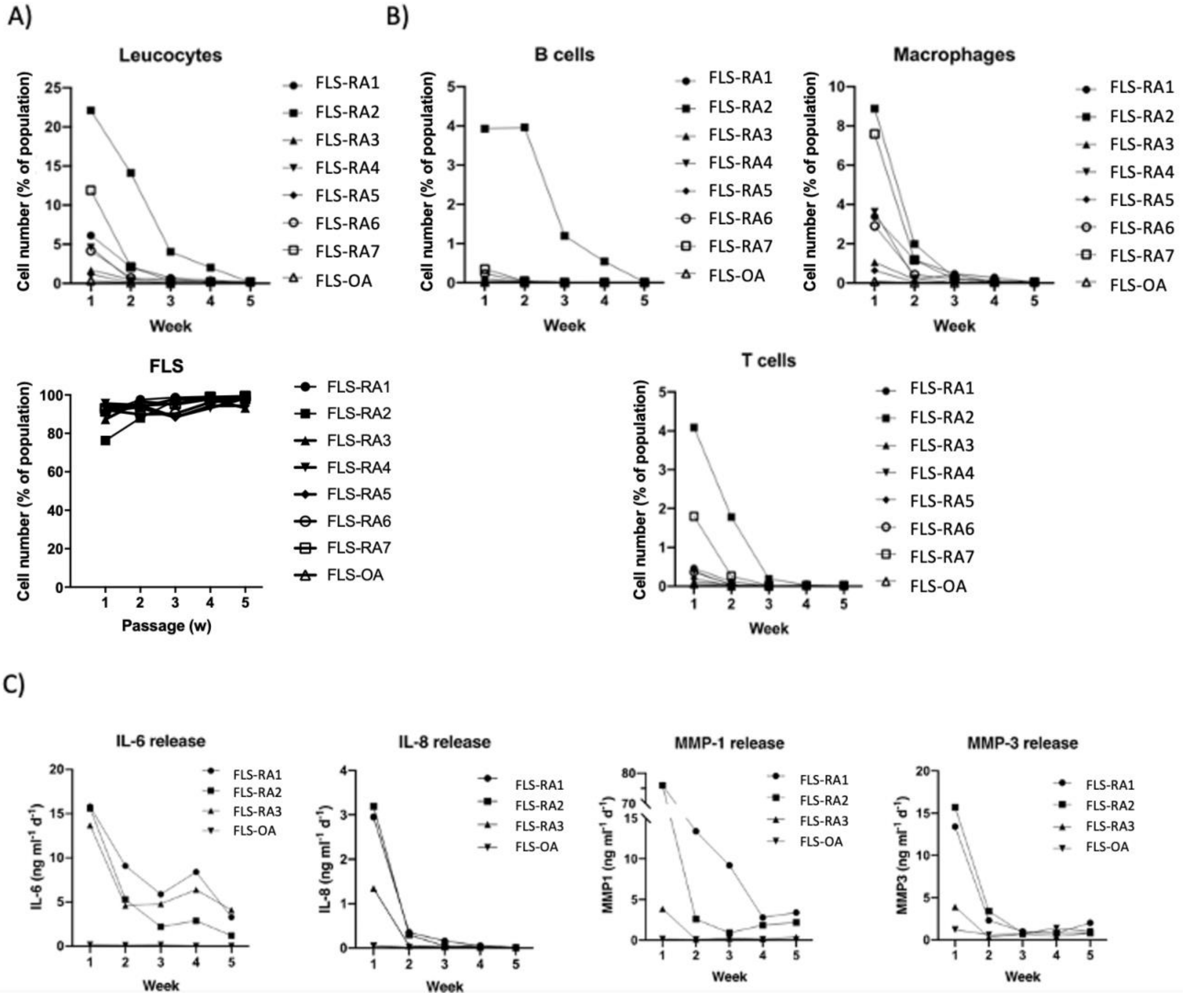


ESI Fig. 3 Characterization of patient-derived cell populations using flow cytometric and cytokine analysis. A,B) Cell distribution for patient-derived fibroblast-like synoviocytes throughout five passages of subcultivation. (RA; Rheumatoid arthritis. OA; osteoarthritis.Data is expressed as % of total gated cell population for n=8 individual patient-derived samples. See ESI Table 1 for information of staining panels.) C) Secretion of proinflammatory cytokines and degradative enzymes throughout five subcultivation passages in 2D culture using Luminex multiplex technology. (RA; Rheumatoid arthritis. OA; osteoarthritis. n=4 individual patient-derived synovial samples)

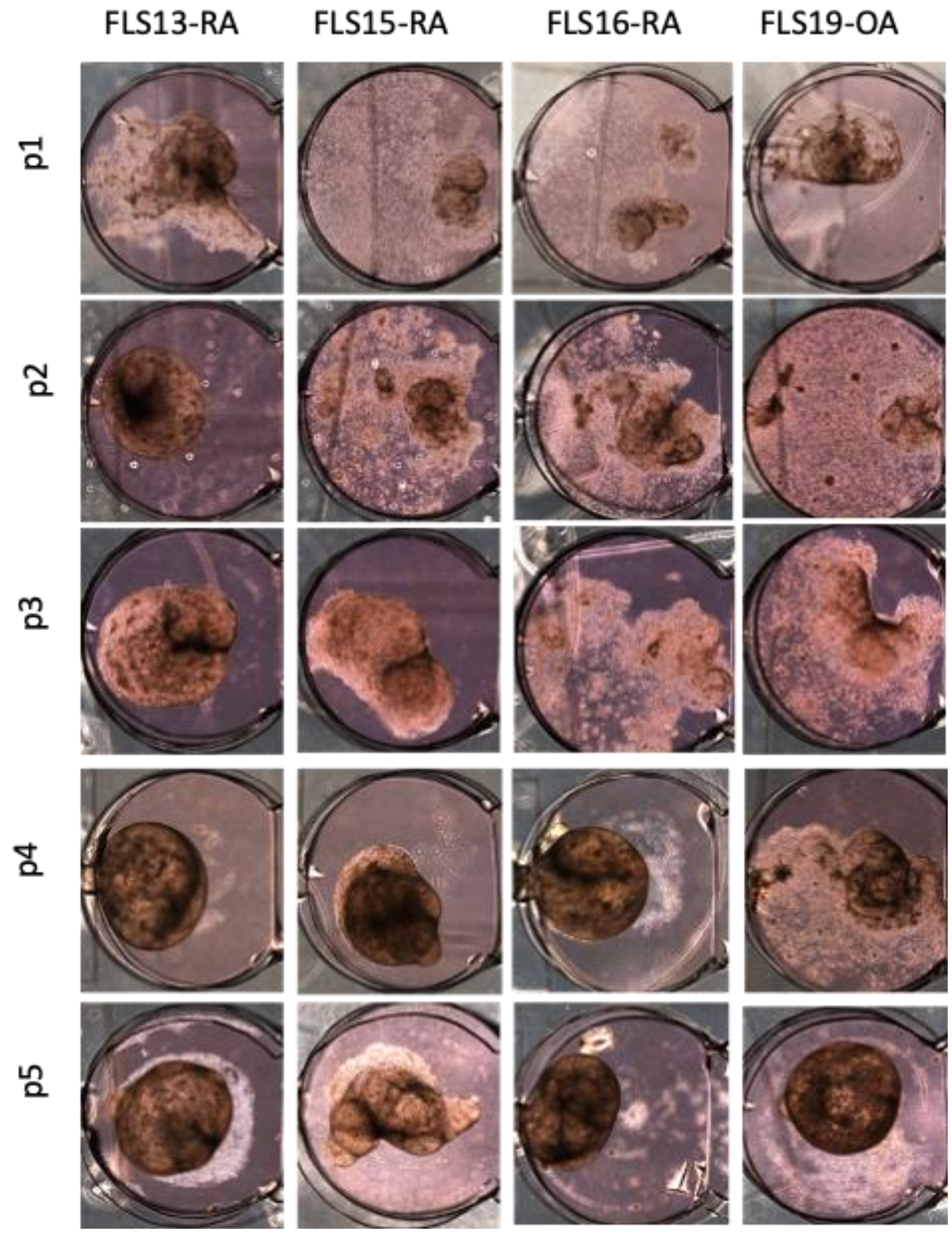

ESI Fig. 4 Representative microscopic images of cultured synovial organoids of different FLS passages up to 5 weeks at day 28 post-seeding.

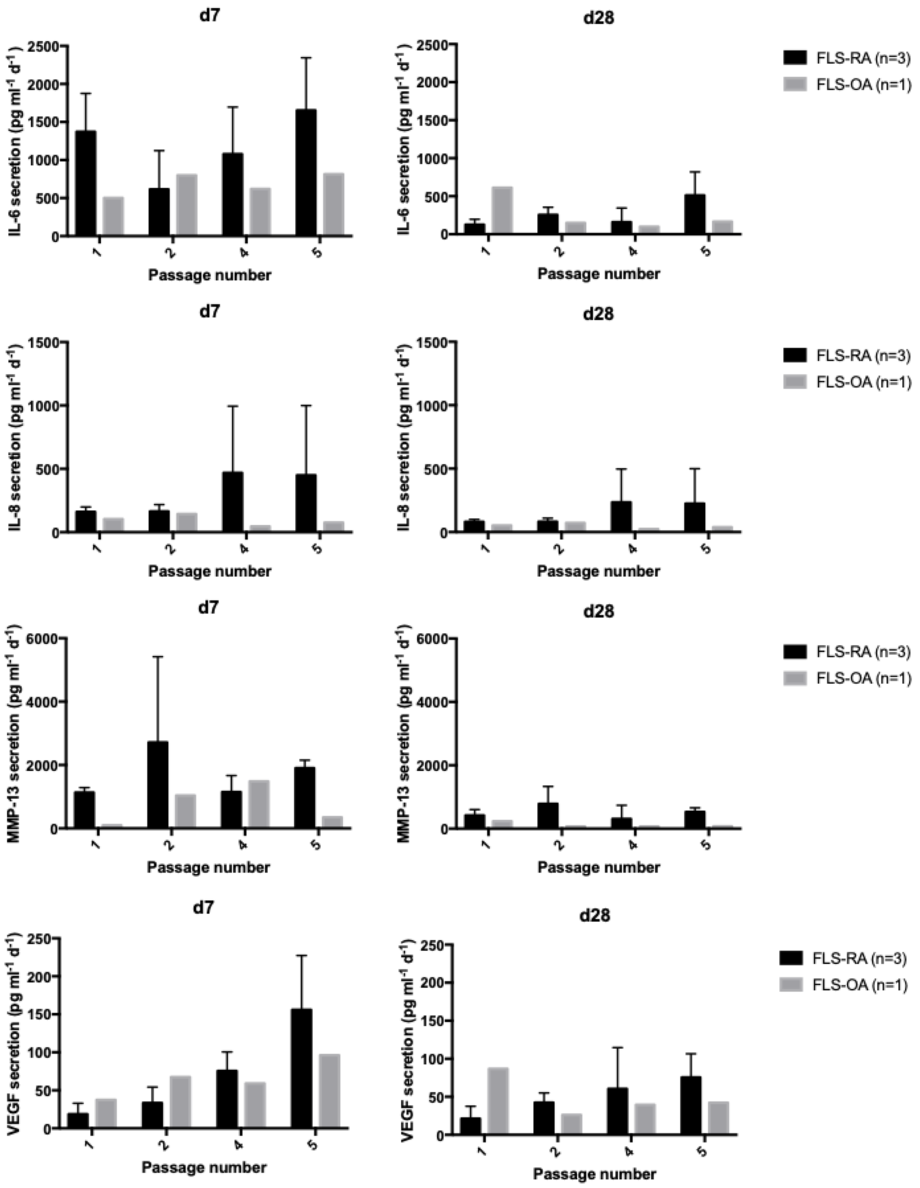


ESI Fig. 5 Secretion levels of proinflammatory molecules including IL-6, IL-8, MMP-13 nd VEGF throughout five subcultivation passages as 3D organoids at day 7 and 28 post-seeding using Luminex multiplex technology. (RA; Rheumatoid arthritis. OA; osteoarthritis. n=1-3 individual patient-derived synovial samples)

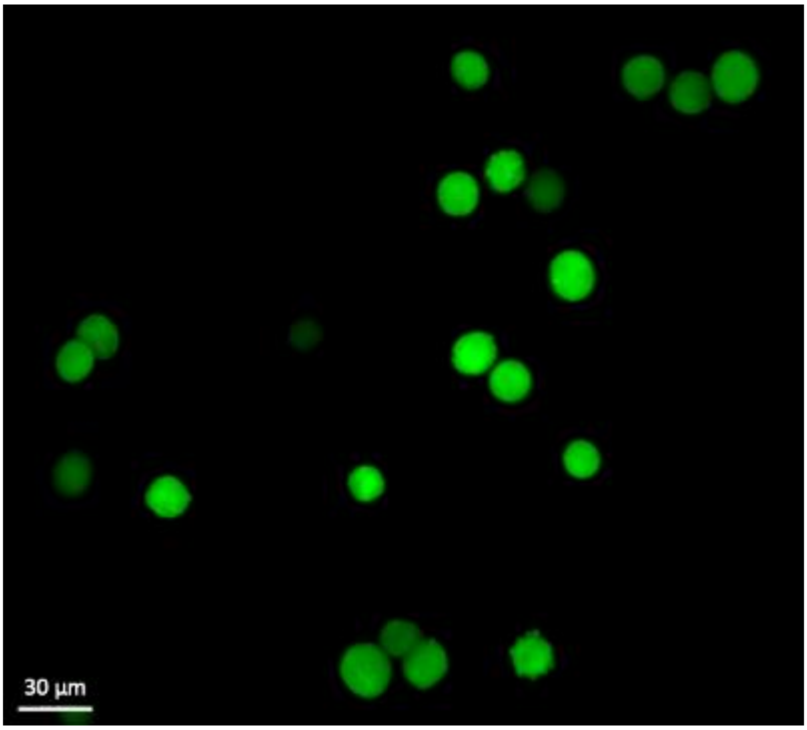

ESI Fig. 6 Fluorescence images of CMFDA-stained primary chondrocytes in the chondral Fibrin-based construct.

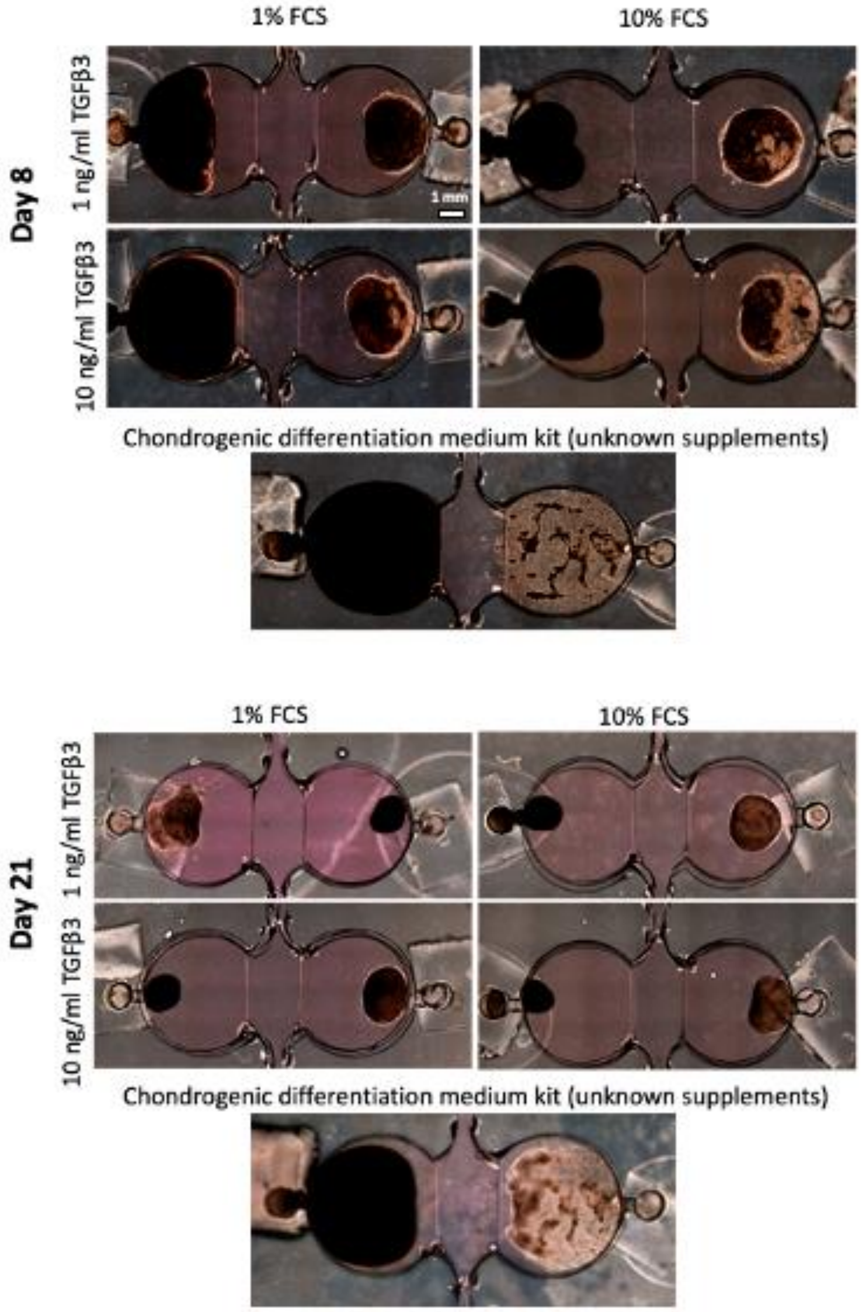


ESI Fig. 7 Representative microscopic images of co-cultured chondro-synovial organoids for different culture medium types and growth factor supplements over 21 days of culture (left chamber: chondral organoids; right chamber: RA-FLS organoids)

### Author contributions

A.F., E.I.R., I.O.C., M.R., R.A.B., and S.S. performed the experiments and analysed the data. F. S. and J. H. provided primary patient tissue. S.S., B.B., W.H. and H.R. provided expertise and assistance with fibrin-based organoids. H. P. K., M.R. and P. E. supervised the work. M.R., P.E., H.P.K., R.W. and S.T. edited the manuscript.

### Conflicts of interest

There are no conflicts to declare.

### Acknowledgements

This project was supported by the Austrian Federal Ministry of Education, Science and Research (BMBWF; Grant agreement number 1612889) and the Vienna Science and Technology Fund (WWTF, project number LS13-092).

### Notes and references

† These authors contributed equally.
